# Contribution and functional connectivity between cerebrum and cerebellum on sub-lexical and lexical-semantic processing of verbs

**DOI:** 10.1101/2023.03.23.533862

**Authors:** Azalea Reyes-Aguilar, Giovanna Licea-Haquet, Brenda I. Arce, Magda Giordano

**Affiliations:** Department of Psychobiology and Neuroscience, Faculty of Psychology, Universidad Nacional Autónoma de México, CDMX, México; Department of Behavioral and Cognitive Neurobiology, Instituto de Neurobiología, Universidad Nacional Autónoma de México, Querétaro, México

**Keywords:** lexical-semantic level, abstract-concrete concepts, grounded cognition, cerebellum

## Abstract

Language comprehension requires sub-lexical (e.g., phonological) and lexical-semantic processing. We designed a task to compare the sub-lexical and lexical-semantic processing of verbs during functional magnetic resonance imaging (fMRI). Likewise, we were interested in the dichotomous representation of concrete-motor versus abstract-non-motor concepts, so two semantic categories of verbs were included: motor and mental. The findings support the involvement of the left dorsal stream of the perisylvian network for sub-lexical processing during the reading of pseudo-verbs and the ventral stream for lexical-semantic representation during the reading of verbs. According to the embodied or grounded cognition approach, modality-specific mechanisms, i.e.,, sensory-motor systems, and the well-established multimodal left perisylvian network contribute to semantic representation for concrete and abstract verbs. The present study detected a preferential modality-specific system for abstract-mental verbs. The visual system was recruited by mental verbs and showed functional connectivity with the right crus I/lobule VI from the cerebellum, suggesting the existence of this network to support the semantic representation of abstract concepts. These results confirm the dissociation between sub-lexical and lexical-semantic processing and provide evidence about the neurobiological basis of semantic representations for abstract verbs.

## Introduction

Language comprehension of spoken words requires sub-lexical (e.g., phonological) and lexical-semantic processing (1–3). According to the dual stream model, the neurobiological substrate in charge of these processes involves two networks in the left perisylvian brain regions (3,4). The first network is a dorsal stream, strongly left-dominant, integrated by structures in the posterior and ventral frontal lobe, posterior temporal lobe, and parietal operculum, involved in translating acoustic speech signals into articulatory representations, essential for speech development and normal speech production (3). The second less understood network is the ventral stream, bilaterally organized and integrated by structures in the anterior and posterior temporal lobe, involved in processing speech signals by mapping phonological representations onto lexical-semantic representations, sound to meaning mapping (1).

For reading, according to the dual-route model of visual word recognition of Coltheart (2005), there are two different routes, a direct/lexical for direct recognition of whole words based on their stored representations requiring the perisylvian ventral stream mechanism (1,6), and an indirect or non-lexical route which involves the assembly of words from individual letters (i.e., the grapheme) and sound (i.e., phoneme) as a function of the dorsal stream (1). The non-lexical route is involved in reading new or unknown words or pseudo-words, i.e., pronounceable letter strings (e.g., RAME), for which there is no lexical entry. For pseudo-words, the perceptual analysis must be reconstructed letter-by-letter during listening or reading to build a word-form representation. However, reading involves additional visual resources for word-form representation and access to the mental lexicon (7,8). Previous studies have found that word reading involved a left language network: superior temporal gyrus/sulcus, middle temporal gyrus, angular gyrus, inferior frontal gyrus, and middle frontal gyrus, whereas pseudo-word reading produced activation in an attentional network that included anterior/posterior cingulate, parietal cortex (7,9), and left ventral frontal regions (10). Another study found that pseudo-words elicited more activation than words in the left dorsal stream, including the supramarginal gyrus in the temporoparietal cortex, precentral gyrus, and insula (11). Thus, all of these brain regions may be part of the non-lexical dorsal route, considered in the dual-route model (5,12).

Moreover, previous studies have shown that language processing in the brain involves left perisylvian regions and extends into sensory-motor and subcortical areas such as the thalamus and cerebellum (10,13–16). It has been shown that motor brain systems contribute to comprehending concrete and abstract concepts, and their recruitment may be flexible as a function of contextual semantic representations (17–20). According to the modality-specific theory, these brain regions may be involved in semantic representation because this representation results from the rehearsal of sensory-motor experiences related to the specific meaning that words carry with them (21). Thus, the modal systems would work together in the semantic processing of both concrete words and some abstract words with activation of consolidated sensory-motor experience (8,17–20). With this evidence, the embodied and grounded cognition approach offers an alternative by considering that conceptual representation is grounded in sensory-motor processes associated with specific contexts and situations (22).

On the other hand, “amodal” theories claim that semantic processing recruits multimodal and integrating brain regions since sensory-motor formats become lost or detached during information integration (23). For example, according to modality-specific theories, the semantic representation of verbs such as ‘write’ involves sensory-motor systems related to the action of moving, the motor experience of the most dexterous hand (19,24), while amodal theories propose that semantic integration mechanisms occur in multimodal regions such as the left perisylvian regions (25).

Although most studies of semantic representation have been done with nouns, recently, verbs have been studied concerning semantic and syntactic representation. Verbs, compared to nouns, are associated more with the recruitment of frontal regions (26,27). Thus, as a fundamental lexical class, verbs denote concrete-motor movement (e.g., ‘write’) or abstract-mental actions (e.g., ‘think’). The involvement of motor and pre-motor brain regions has been shown during the reading of verbs denoting actions (25,28,29). A functional magnetic resonance imaging (fMRI) study with verbs in Spanish (30) compared concrete and abstract verbs relative to pseudo-verbs using a region of interest (ROI) approach on frontal and temporal regions. The authors found that activation for concrete verbs was located on frontal motor regions, while the activation for abstract verbs was anterior to frontal motor regions. In the same way, Dalla Volta et al. (2014) detected more activation in the right somatosensorial regions and the left parietal lobe for concrete verbs, while abstract verbs evoked higher activation in the prefrontal cortex outside of motor areas. These results support the hypothesis that abstract concepts recruit multimodal brain regions, while concrete concepts recruit sensory-motor regions.

Conversely, recent studies have demonstrated that abstract concepts have associated motor and perceptual properties. Their semantic representation includes the braińs modal systems for perception, action, and introspection (31), supporting the embodied and grounded cognition models (17,18,20). The involvement of modal systems in the semantic representation of abstract verbs could depend on the context or strategies, for instance, making use of visual experiences (8,22). In this regard, visual brain regions are relevant for processing some abstract verbs (mental verbs such as ‘understand’) that afford visual information (18,21). On the other hand, cortical and subcortical motor regions should be involved in semantic representations of motor movements, e.g., motor verbs such as ‘write’, as well as regions related to lateralized dominance for behavioral motor proficiency, motor experience and handedness (17,22).

Despite the recent increase in the number of studies about verbs, the neural basis of sub-lexical and lexical-semantic processing of verbs needs to be better understood. In this study, we focused on verbs as a lexical category that can represent concrete or abstract actions. To compare the contribution and functional connectivity of brain regions on sub-lexical and lexical-semantic processing of verbs, in addition to verbs, we included pseudo-verbs and strings of letters with the characteristic ending part of the verb in the present tense in Spanish (i.e., *ar, er, ir*). Thus, we contrasted verbs and pseudo-verbs, which we expected would confirm the recruitment of the dorsal stream for pseudo-verbs and the ventral stream for verbs. Likewise, we were interested in the dichotomous brain representation of concrete-motor versus abstract-non-motor action verbs because, as mentioned above, the brain correlates are controversial (17,22,25,32). Thus, we evaluated the involvement of integrating multimodal, and modality-specific brain regions in the processing of motor verbs (with concrete meaning) versus mental verbs (with abstract meaning). According to the embodied or grounded cognition approach, we expected to find that both multimodal and modality-specific cortical regions would be involved in processing both categories of verbs (17,33). However, we expected differences in modality-specific regions between verb categories, motor verbs would engage the sensory-motor areas, and the mental verbs, the visual brain areas.

## Methods

### Participants

Twenty-four volunteers (12 women) between 21 and 35 years old (M = 26.75, SD = 3.95), all native-Spanish speakers, participated in this study. Sample size was calculated to detect a within-subject effect of a large size (*d* = 0.8), with 80% power, and a low alpha value (0.01) using a two-tailed one-sample *t-*test. Participants showed no neurological or psychiatric disorders according to the Mexican version of Symptom Checklist 90 (34). After the participants were informed of the study procedures and confidentiality, they signed the written informed consent to participate in the experiment. The experimental protocol was approved by the Ethics in Research Committee of the Instituto de Neurobiologia-UNAM in compliance with federal guidelines of the Mexican Health Department (NOM-012-SSA3-2012), which agree with international regulations.

### Stimuli

We selected the verbs in Spanish according to two categories, motor, and mental verbs, based on their psycholinguistic properties from the ADESSE database (http://adesse.uvigo.es/). Firstly, for motor verbs, we selected verbs from the material class related to movements in space and modification of objects. For mental verbs, as a second category, we selected those that belong to the mental class associated with perception, cognition, and choice (*elección* in Spanish). A total of 330 verbs resulted from this procedure: 167 motor verbs and 163 mental verbs. Then, the frequency of use for each verb was obtained in Sketch-Engine (35), http://www.sketchengine.eu) and LEXMEX-Spanish (36) corpora if one verb was not in both corpora, it was excluded. Following this procedure seven verbs were excluded. Then, both verb categories were matched according to their frequency of use in the LEXMEX-Spanish corpus. In total, 139 mental verbs and 151 motor verbs were selected. Then, the verbs were checked, and both categories were matched according to their number of syllables; two were excluded (i.e., they had four or more syllables). Finally, 288 verbs were selected, 139 mental verbs and 149 motor verbs.

These 288 verbs were presented to thirty young participants (an independent sample from that of the fMRI study), native-Spanish speakers, for reading and rating each verb on five psycholinguistic properties (Likert scale from 1 (less) to 6 (more)): motor-mental relatedness, concreteness-abstraction, imageability, emotional valence, and arousal. In four out of five psycholinguistic properties, there were differences between both categories according to a Wilcoxon Signed-Rank test. Motor verbs were classified as motor while mental verbs as mental (Z = 162, p < 0.001); in the concreteness-abstraction scale, motor verbs obtained higher concreteness value while mental verbs as more abstract (Z = 247, p < 0.001); motor verbs were rated with more imageability than mental verbs (Z = 18340, p < 0.001); and mental verbs showed higher scores in arousal respect to motor verbs (Z = 8323, p < 0.01). There were no differences in emotional valence between verb categories. We selected those motor verbs with higher motor relatedness, i.e., scores < 3, and mental verbs with high mental relatedness, i.e., scores > 4, without overlap between categories. Similarly, there was no overlap between categories on concreteness-abstraction, motor verbs had scores < 3, and mental verbs, > 3; or on imageability, motor verbs scored = 6, and mental verbs < 6. As a result, we got two verb categories with similar frequency of use and number of syllables, but different degrees of motor-mental relatedness, concreteness-abstraction, and imageability, i.e., 112 motor-verbs, e.g., *caminar* [walk], and 112 mental-verbs, e.g., *entender* [understand]. Then, we built pseudo-verbs with a comparable level of pre-lexical familiarity. For each verb, a pseudo-word was created, changing the position of consonants but maintaining the vowels and the ending [ar, er, ir], which is a lexical indicator of the verbs in Spanish, e.g., for verb = *caminar*, the pseudo-verb = *nacimar*, conserving the phonological (i.e., sub-syllabic) structure of the words; all resulting letter sequences were pronounceable and corroborated in a dictionary in Spanish (*Real Academía Española*, https://dle.rae.es/nacimar?m=form) if any of them was identified with a meaning, that letter sequence was eliminated. After this, 112 pseudo-verbs were randomly selected for the fMRI study. Finally, one last category of stimuli was included. A string of symbols was built for each verb, and a pair of the same symbol or characters was included for each syllable of the verb (e.g., for *caminar*, the stream of symbols was $$##&&). After this, half of those symbol strings were randomly selected to obtain 112.

### Identification task

Four categories were included: mental verbs, motor, pseudo-verbs, and symbols. The stimuli were presented for 1000 ms with 2000 ms interstimulus intervals. Ten stimuli from one category were shown in a block, and three were repeated randomly to create the “one-back” detection task. Blocks were separated by 12-sec blank intervals (Fig 1). Two blocks of each stimulus category were presented pseudo-randomly in each of the eight runs. Each run lasted approximately 6 min. Participants were instructed to identify the current stimulus and indicate during the stimulus presentation if that stimulus was the same as the previous one (one-back detection task). The stimuli were presented on a gray background using the PsychoPy® software (37,38), and a projection system consisting of MR-compatible goggles from NordicNeuroLab (Bergen, Norway). Additionally, a button system from NordicNeuroLab was used to record the participants’ responses.

**Fig 1.**
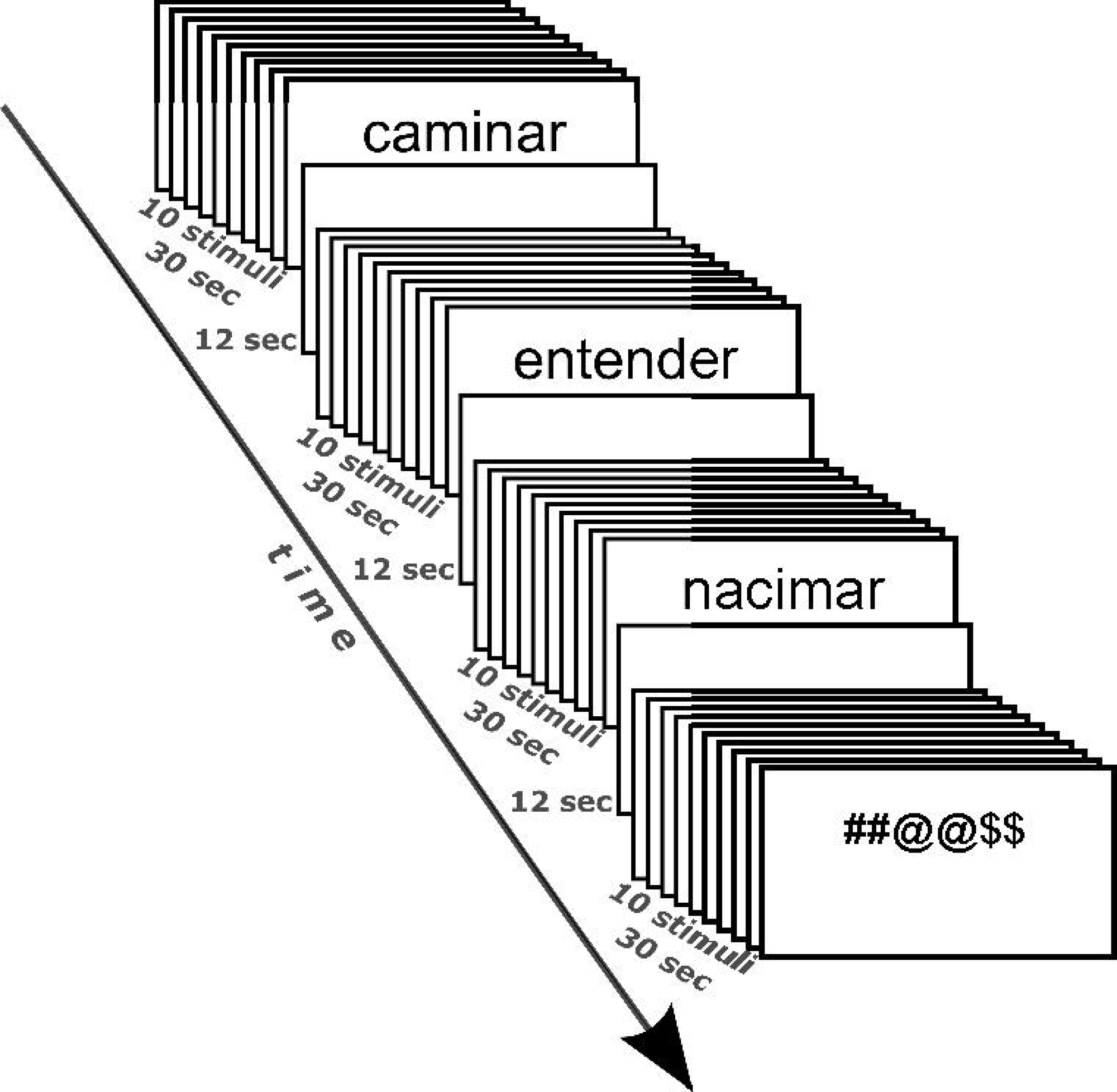
The stimuli were presented for 1000 ms with 2000 ms inter-stimulus intervals. Each block included ten stimuli from one category, and subjects performed a one-back detection task;12-sec blank intervals separated blocks. Two blocks of each stimulus category were presented pseudo-randomly in each of the eight runs.

### Procedure

#### fMRI study

As indicated before, participants were instructed to identify the current stimulus (i.e., verbs and pseudo-verbs, and symbols strings) and indicate during the stimulus presentation if that stimulus was the same as the previous one (one-back detection task). Regardless of stimulus type, reading or identifying stimuli engages similar visual processing, but only verbs and pseudo-verbs require phonological processing, and only verbs involve lexical-semantic processing. Thus, we expected motor and mental verbs to involve differential semantic processing because they varied in motor-mental relatedness, concreteness-abstraction, and imageability.

#### Behavioral testing

On a different day, participants returned to complete the vocabulary subscale of the Wechsler Adult Intelligence Scale (WAIS-IV, (39)) as an index of lexical performance for each participant and three varieties of verbal fluency tasks. For one minute, they had to produce certain types of words: verbs as a lexical category (verbs fluency), animal names (semantic fluency), and words with “m” as the first letter (phonological fluency). To avoid a fatigue effect, the order of behavioral tasks was counterbalanced at the subject level. Finally, the Edinburgh Handedness Inventory (EHI; 40) was applied to assess the dominance of the participant’s right or left hand in everyday motor activities (M = 76.04, SD = 20.91).

#### Imaging data acquisition

fMRI imaging was performed on a 3.0T GE MR750 scanner (General Electric, Waukesha, WI) using a 32-channel head coil. Functional imaging included 38 slices, acquired using a T2*-weighted echo-planar imaging sequence with TR/TE 2000/40 ms, field of view of 25.6 cm, a 64 × 64 matrix, and 4-mm slice thickness, resulting in a 4 × 4 × 4 mm3 isometric voxel. High-resolution structural 3D-T1-weighted images were acquired for anatomical localization. These images were acquired with 3D spoiled gradient recall (SPGR), resolution of 1 × 1 × 1 mm3, TR = 8.1 ms, TE = 3.2 ms, flip angle = 12°, inversion time = 0.45, covering the whole brain.

#### fMRI data analysis

For quality control of the BOLD data, following a maximum absolute motion of more than 2mm as an exclusion rule, one of eight runs was excluded for three subjects. MRI data were analyzed using FSL (FMRIB’s Software Library, www.fmrib.ox.ac.uk/fsl)(41). Statistical analysis was performed with FMRI Expert Analysis Tool using FMRIB’s Improved Linear Model (FEAT FILM) Version 6.0.1. Each participant’s data were brain extracted, motion and slice timing corrected, and normalized onto MNI common brain space (Montreal Neurological Institute, EPI Template, voxel size 2mm x 2mm x 2mm). Data were smoothed using a gaussian filter (full-width half maximum = 6mm) and high-pass filtered during analysis. Blood oxygen level-dependent (BOLD) signal was examined during the stimuli presentation when participants were instructed to read or identify the stimulus (verb, pseudo-verb, or symbols). Statistical analysis of event-related hemodynamic changes was carried out per the general linear model (GLM, Friston et al., 1995). The model included the following regressors: motor verbs, mental verbs, pseudo-verbs, and symbols. First-level fMRI analysis data was performed to identify regions that increased BOLD signal intensity for each of the four categories of stimuli relative to blocks of blank intervals for each run with a significance threshold criterion of *Z* > 2.3. Since each subject responded to the experimental paradigm in eight independent runs, to estimate a map of the brain regions involved during the identification of each stimuli category, a mid-level analysis was carried out using a fixed-effects model, which averaged the activity of verbs and pseudo-verbs during each of the eight runs respect to symbols: a conjunction analysis was done with both verbs categories, motor-verbs ∩ mental-verbs > symbols contrast, and pseudo-verbs > symbols contrasts. To test whether verbs recruited different mechanisms relative to pseudo-verbs, a conjunction analysis was done with both verb categories: motor-verbs ∩ mental-verbs > pseudo-verbs, and pseudo-verbs > motor-verbs ∩ mental-verbs contrasts. Then, we compared verb categories: motor > mental and mental > motor contrasts. Finally, we conducted this analysis with the removal of mechanisms related to visual processing (symbols) and phonological processing (pseudo-verbs): [motor > symbols] > [mental > symbols], [mental > symbols] > [motor > symbols], [motor > pseudo] > [mental > pseudo], and [mental > pseudo] > [motor > pseudo] contrasts. To identify activations at the group level, we used a third-level analysis using FLAME 1 (FMRIB’s Local Analysis of Mixed Effects) with a cluster significance threshold criterion of *Z* > 2.3 with *p* < 0.05 corrected for multiple comparisons with Gaussian Random Field (GRF) for results at the whole-brain level (43).

#### Region of interest (ROI) analysis

ROI analysis was performed on ten regions detected as maxima from an automated meta-analysis using “verbs” as a term in neurosynth (http://neurosynth.org/, (44). For the voxel as a maxima activation, an 8-mm *spherical* ROI was built for each ROI. Nine ROIs were localized on the cerebral cortex (seven on the left hemisphere, two on the right hemisphere according to Harvard-Oxford Cortical Structural Atlas), and the last one, on the right cerebellum (Cerebellar Atlas in MNI152 space). The ROIs were located as follows: left temporal occipital fusiform cortex (L-FusOcc [−40, −50, −20]), left inferior frontal gyrus (L-IFG [-50, 16, 16], pars opercularis), left superior lateral occipital cortex (L-LOC [-26, −60, 48], superior parietal lobule and angular gyrus), left anterior and posterior middle temporal gyrus (L-MTG-ant [-60, −50, 0], and L-MTG [-58, −10, −14], respectively), left supplementary motor area (L-SMA [-2, 8, 60]), left supramarginal gyrus (L-SMG [-60, −26, 28]), right orbitofrontal cortex (R-orb [36, 26, −4]), right superior temporal gyrus (R-STG [56, −32, 2]), and right cerebellum (R-Cerebellum [34, −64. −29], crus 1 and lobule VI). Then, for each ROI, the average percent signal change from calculated semantic contrasts in the GLM (mental vs. motor verbs) was calculated. Finally, a correlation analysis was carried out with these values and the scores obtained from the behavioral tests.

#### Psychophysiological interaction analysis

To test the functional connectivity modulated by phonological and lexical-semantic processing, psychophysiological (PPI) analysis was conducted with those seeds ROIs that showed a correlation between semantic contrast in the GLM analysis and scores from behavioral tests (i.e., L-SMG, L-LOC, L-MTG, L-SMA, and R-Cerebellum). In addition, we conducted a whole-brain PPI analysis for each seed to know how each one of the seeds (time series of the brain region, i.e., seed, as physiological regressor) was coupled with other brain regions during the reading of verbs as compared to symbols and pseudo-verbs (psychological regressors).

Each ROI was projected on the pre-processed functional images, i.e., eight runs for each participant. Then, the time series of BOLD activity was extracted using fslmeans utility as an average across all voxels within each seed ROI for each data set. Finally, the PPI analysis was conducted for every ROI separately using FEAT (FMRI Expert Analysis Tool) Version 6.00, part of FSL (FMRIB’s Software Library www.fmrib.ox.ac.uk/fsl).

First-level GLM analysis included nine regressors: a physiological regressor (i.e., time series of the seed), four psychological regressors corresponding to the four stimulus categories (based on the design of the one-back detection task), and the last four regressors for interaction between psychological and each physiological regressor (PPI). Psychophysiological interaction was determined by testing for a positive slope of the PPI regressor. Individual contrast images for PPI analysis were the same as whole brain analysis and were entered into the subject-level analysis. Data for PPI analysis, as well as GLM, were processed with FEAT, part of FSL. Subject-level analyses were performed separately for each condition. Finally, the contrast images generated at the subject-level analyses were entered into the group-level analyses.

#### Statistical analysis of behavioral measures

Behavioral data were analyzed using R 3.6.1. We compared the performance among three verbal fluency tasks with repeated-measures ANOVA. Pearson analysis correlation was used to test whether the scores of behavioral tasks were related. Pearson correlations or partial correlations were calculated to test the relation between scores on behavioral tasks and changes in BOLD signal in ten seeds from those contrasts between categories of verbs.

## Results

### Behavioral measures - linguistic performance

A higher fluency of verbs (F (2, 46) = 33.66, p < 0.001) and semantic fluency (p < 0.001) relative to phonological fluency was detected. No difference was found between semantic and verb fluency.

The vocabulary task was related to verb fluency (r = 0.65, p < 0.001, Spearman, FWE corrected) and to semantic fluency (r = 0.52, p < 0.01, Spearman, FWE corrected) but not to phonological fluency. Then, we tested if there was a relation between performance on linguistic tasks and EHI as a measure of lateralized everyday motor behavior. Verb fluency showed a positive correlation with phonological fluency (r = 0.60, p < 0.01, Spearman, FWE corrected) and a negative correlation with EHI (r = −0.43, p < 0.05, Spearman, non-corrected, Fig 2).

**Fig 2.**
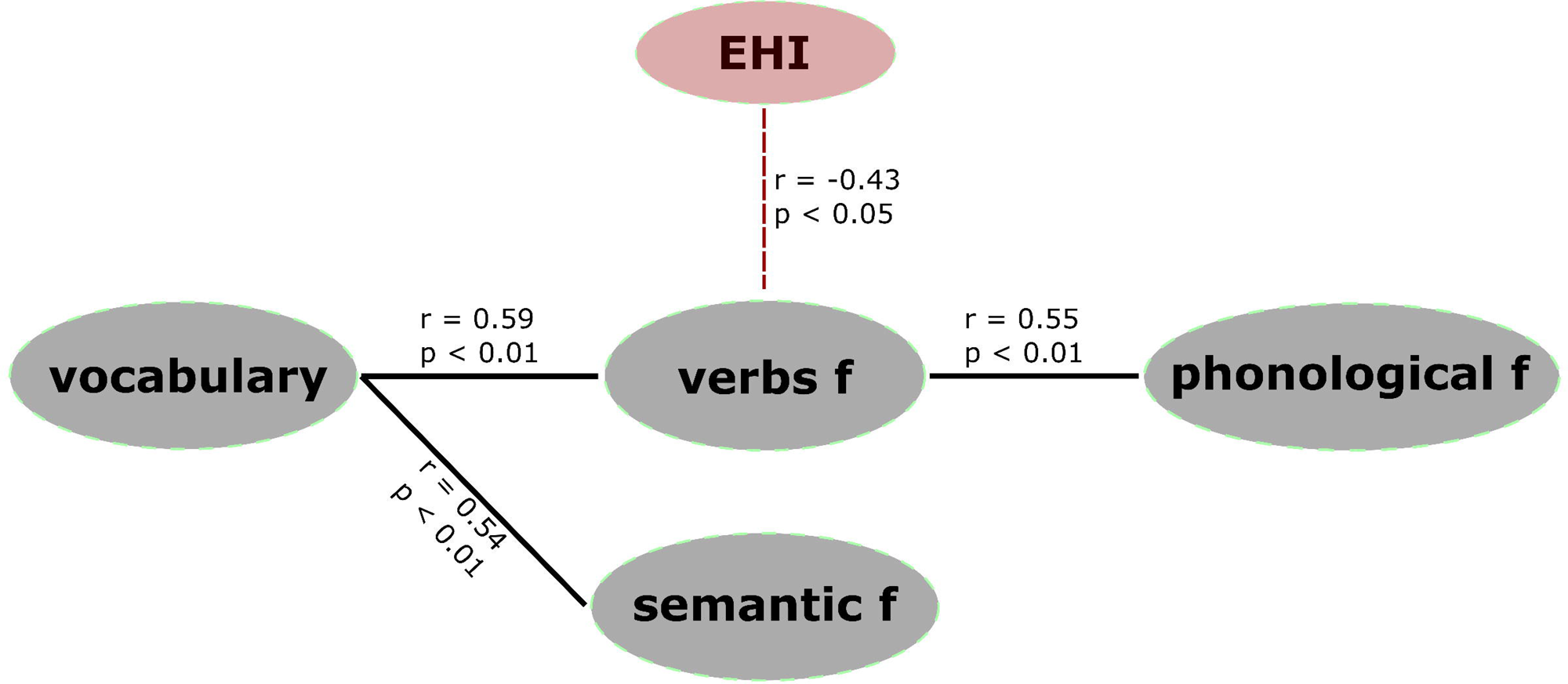
Correlation among scores on behavioral tasks. Vocabulary task of WAIS-IV, verb fluency, semantic fluency, phonological fluency, and EHI (Edinburgh Handedness Inventory), f (fluency).

### fMRI results

In the fMRI study, the participants successfully performed the one-back detection task with an overall 99% of correct responses, which confirmed that the stimuli were attended for visual, phonological, and lexical processing to determine whether two consecutively stimuli were the same or not. Furthermore, there were no differences in the correct responses across stimuli categories, so the differences detected in the BOLD signal were not due to the difficulty of the stimulus categories.

Results from the GLM analysis showed that, compared to symbols, verbs recruited bilateral frontal, temporal, and parietal cortex (Fig 3, S1 Table). In contrast, pseudo-verbs showed involvement of the left hemisphere language network extending into motor areas such as pre and postcentral gyrus, SMA, and right cerebellum (crus I, crus II, lobules VI, and VIIIa; Fig 3 and, S1 Table).

**Fig 3.**
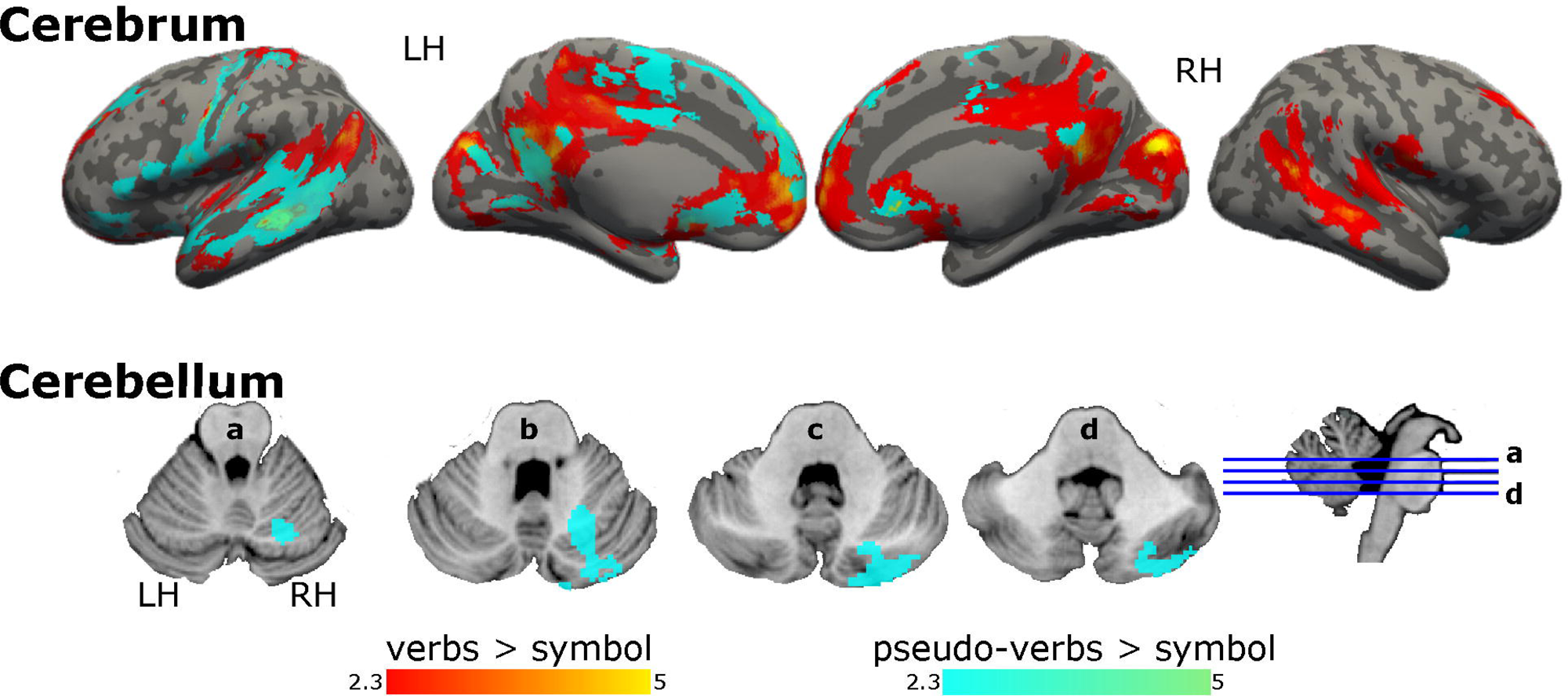
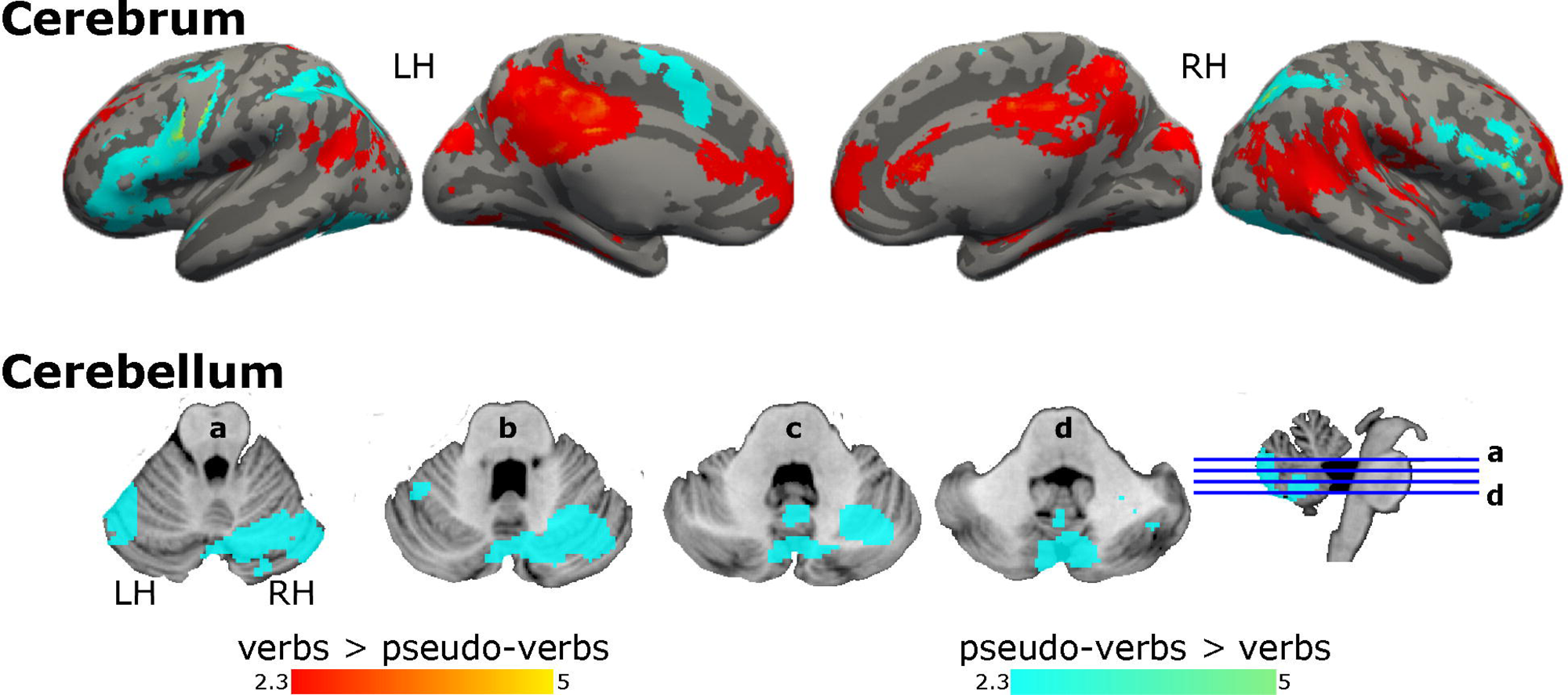
Activation maps for verbs and pseudo-verbs with respect to symbols. Graphical representation of GLM’s results, brain regions activated in the contrast verbs > symbols in red, and pseudo-verbs > symbols in aqua. Coordinate z of multislices in cerebellum: a = −24, b = −29, c = −34, and d = −39. Color bars show z scores. LH: left hemisphere; RH: right hemisphere.

Verbs, in contrast to pseudo-verbs, recruited left posterior temporal, right middle temporal, ventral parietal, and frontal regions with more extension on the right hemisphere, in addition to medial regions: anterior and posterior cingulate cortex, and occipital areas, i.e., lingual gyrus (Fig 4, S2 Table). Moreover, the opposite contrast, pseudo-verbs as compared to verbs, recruited more left-lateralized frontal motor regions, pre and postcentral gyrus, SMA, and right cerebellum (crus I, crus II, lobules VI and VIIb), bilateral superior lateral occipital cortex, and left temporal occipital fusiform cortex including the visual word form area (Fig 4, S2 Table).

**Fig 4.**
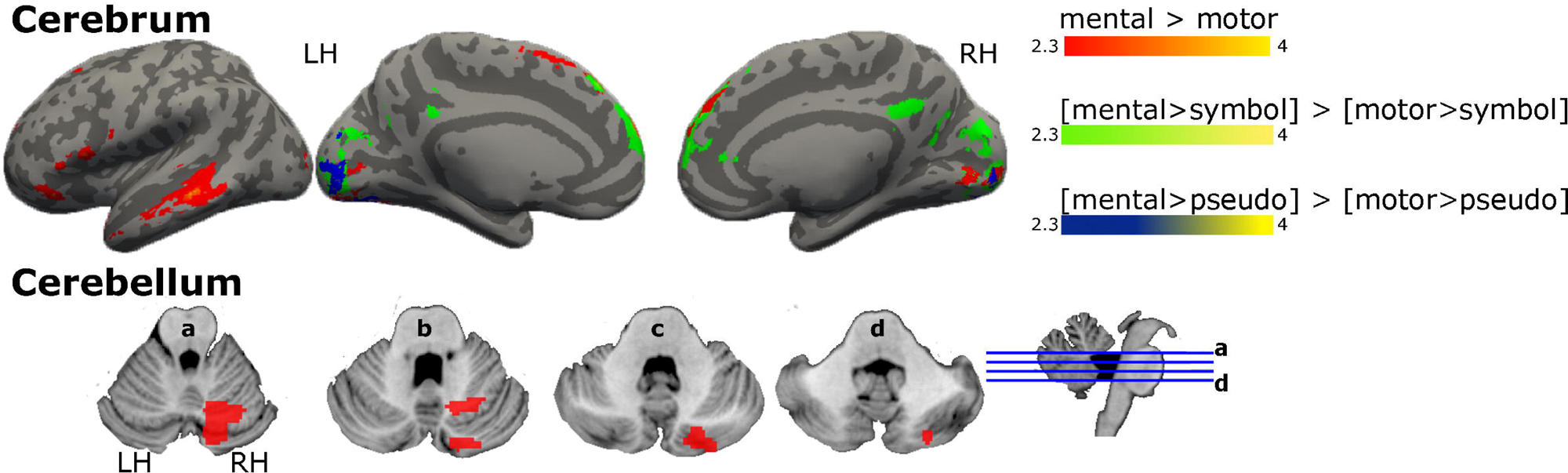
Activation maps for the comparison between verbs and pseudo-verbs. Graphical representation of GLM’s results, brain regions activated in the contrast verbs > pseudo-verbs in red, and pseudo-verbs > verbs in aqua. Coordinate z of multislices in cerebellum:: a = −24, b = −29, c = −34, and d = −39. Color bars show z scores. LH: left hemisphere; RH: right hemisphere.

When comparing the effect of semantic differences between verb categories, we found no effect for the motor > mental contrast. Conversely, the mental > motor contrast evidenced an increase of BOLD signal on the left inferior frontal region, dorsomedial prefrontal cortex, including SMA, left ventral temporal areas and medial occipital regions, and right cerebellum (crus II and lobule VI, Fig 5, S3 Table). Removing the effect of visual processing (i.e., symbols) in the comparison [motor > symbols] > [mental > symbols], resulted in the recruitment of bilateral lateral superior occipital cortex and ventrolateral temporal regions, including the fusiform area (S1 Fig, S4 Table). Whereas the opposite contrast, [mental > symbols] > [motor > symbols] showed an increase of BOLD signal in medial frontal and occipital regions (Fig 5, S3 Table). Finally, when the phonological effect was removed (i.e., pseudo-verbs), the contrast [motor > pseudo] > [mental > pseudo] did not show significant differences in BOLD signal, however the opposite contrast, [mental > pseudo] > [motor > pseudo], recruited the medial occipital areas, and lingual gyrus (Fig 5, S3 Table).

**Fig 5.**
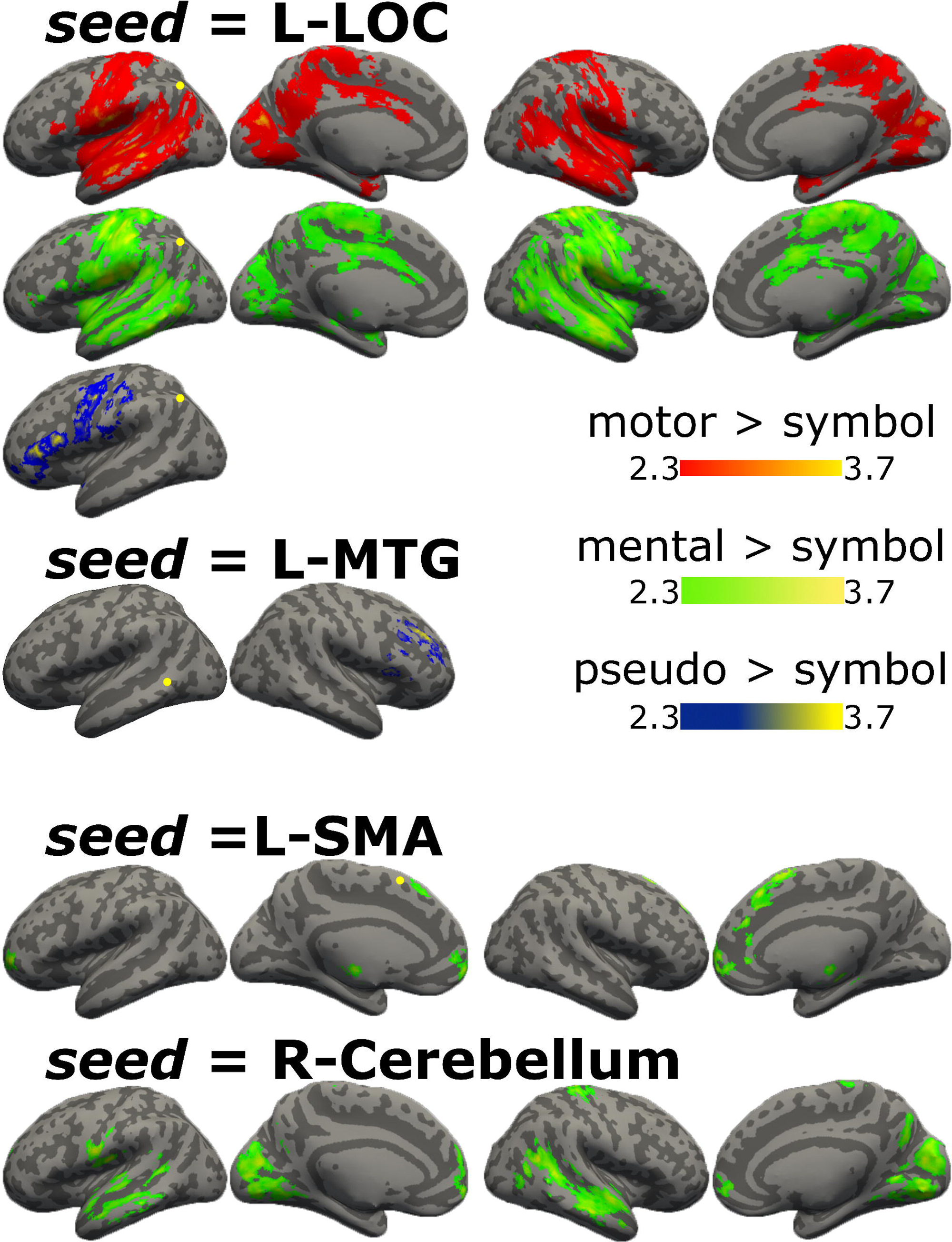
Activation maps for the comparison between mental and motor verbs. Graphical representation of GLM’s results, brain regions activated in the contrast mental > motor verbs in red, same contrast when the effect of visual processing (i.e., symbols) is removed, in green, and when the effect of phonological processing (i.e., pseudo-verbs) is removed, in blue. Coordinate z of multislices in cerebellum: a =: a = −24, b = −29, c = −34, and d = −39. Color bars show z scores. LH: left hemisphere; RH: right hemisphere.

### Region of interest (ROI) analysis

We asked about the relation between behavioral tests and the BOLD signal changes observed in specific language brain areas during the identification task. We defined ten ROIs from a meta-analysis for “verbs” (see details in Methods).

The BOLD signal change for mental > motor verbs contrast in the L-LOC ROI was positively correlated to verbs fluency (r = 0.53, p < 0.01, FWE corrected), and in the L-MTG ROI it was negatively correlated to EHI (r = −0.54, p < 0.01, FWE corrected). Since there was a significant correlation between verbs fluency and EHI, we calculated the partial correlations between EHI and BOLD signal, with verbs fluency as covariable. Negative partial correlations, without correction, were found for motor areas: L-SMA (r = −0.45, p < 0.05) and R-cerebellum (r = −0.42, p < 0.05). Other significant correlations were detected between behavioral measures and BOLD signal change for other contrasts: verbs > symbol and verbs > pseudo-verbs (S5 Table).

### Psychophysiological interaction

Having established that those four ROIs (i.e., L-LOC, L-MTG, L-SMA, and R-Cerebellum) showed a relation to behavioral tasks in the contrast mental > motor verbs, suggesting semantic differences, we conducted a PPI analysis with these four ROIs as seeds. We analyzed functional connectivity of verbs relative to other categories, i.e., symbols and pseudo-verbs, and between categories of verbs, i.e., mental and motor. PPI analysis was calculated for the following contrasts: motor > symbol, mental > symbol, pseudo > symbol, motor > pseudo, mental > pseudo, motor > mental, and mental > motor.

First, we obtained the psychophysiological interaction for motor and mental verbs and pseudo-verbs, with respect to symbols. L-LOC showed functional connectivity with bilateral temporal and parietal regions for motor and mental verbs. While for pseudo-verbs the functional connectivity was only with left regions such as IFG, pre, and postcentral gyrus. L-MTG, as seed, showed functional connectivity with the right dorsolateral prefrontal cortex only for pseudo-verbs. L-SMA and R-Cerebellum were coupled only for mental verbs. L-SMA with bilateral frontal regions and thalamus, while R-Cerebellum with medial occipital regions, bilateral temporal regions, pre and postcentral gyri, and polar frontal regions (Fig 6, S6 Table).

**Fig 6.**
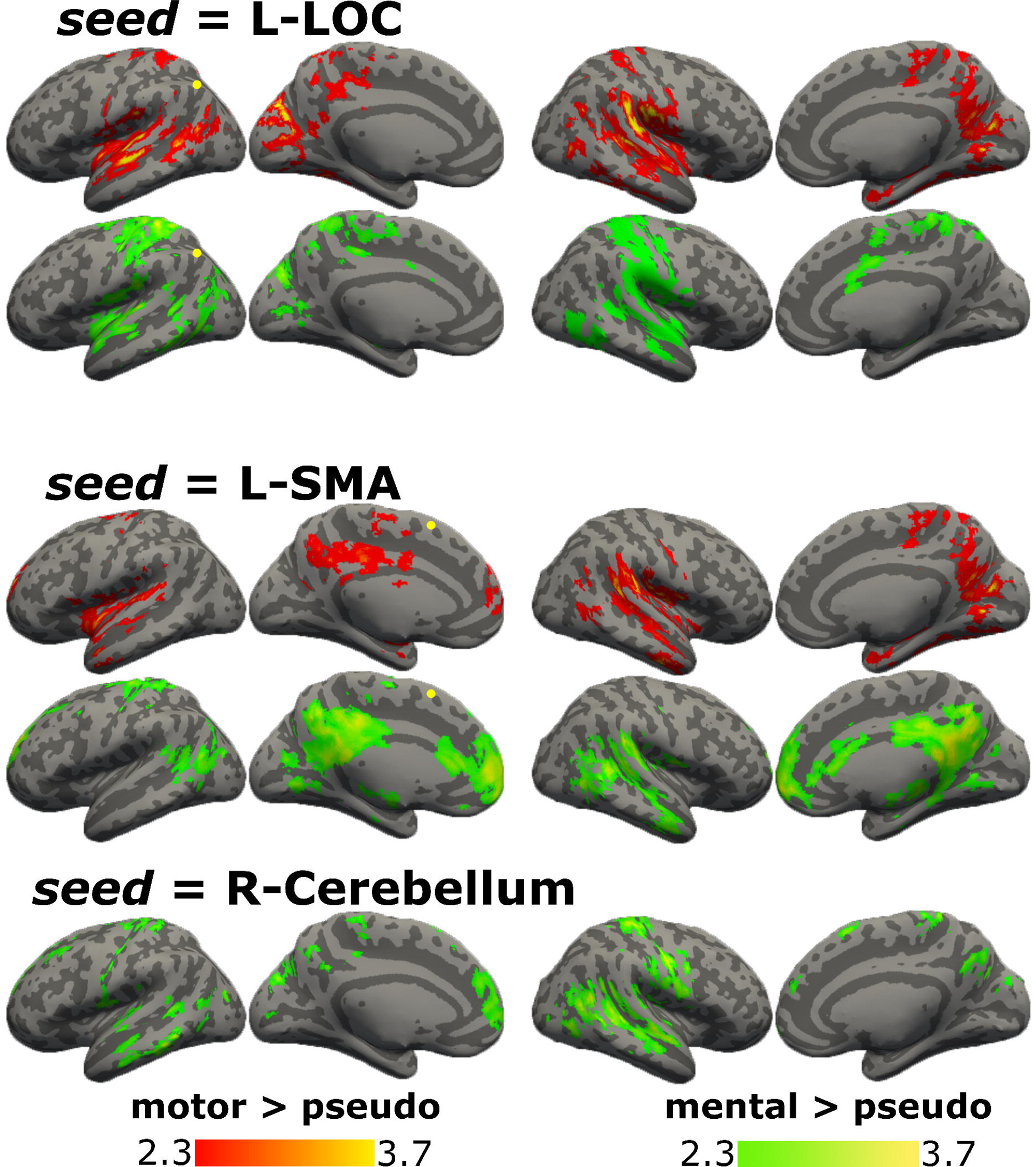
PPI results for motor, mental and pseudo verbs with respect to symbols. Graphical representation of PPI results with seeds in L-LOC, L-MTG, L-SMA, and R-cerebellum, brain regions functionally connected in the contrast motor > symbols in red, mental > symbols in green, and pseudo-verbs > symbols in blue. Color bars show z scores. LH: left hemisphere; RH: right hemisphere.

Then, psychophysiological interaction was calculated for each category of verbs with respect to pseudo-verbs (Fig 7, S7 Table). For motor verbs, L-LOC was connected with bilateral parietal and temporal regions, and for mental verbs, with more extension in postcentral areas. L-SMA, for motor verbs, was connected with bilateral superior temporal regions, posterior cingulate cortex, and frontal pole, while for mental verbs, with posterior cingulate cortex, medial prefrontal cortex, left dorsal parietal regions, and bilateral posterior temporal areas. R-Cerebellum was connected with other regions only for mental verbs: bilateral temporal regions, post and precentral gyri, frontal areas, and precuneus.

**Fig 7.**
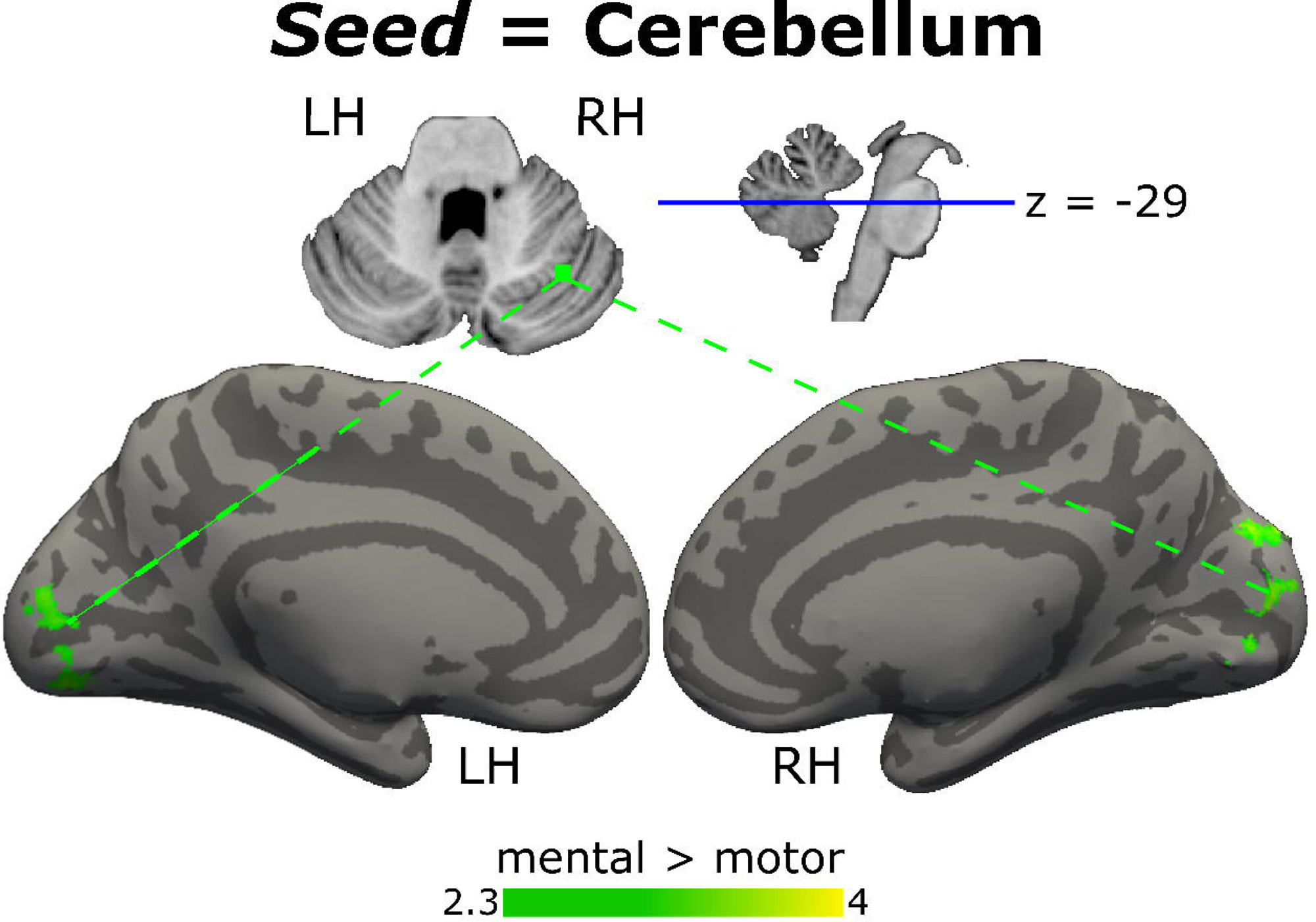
PPI results for motor and mental verbs with respect to pseudo-verbs. Graphical representation of PPI results with seeds in L-LOC, L-SMA, and R-cerebellum, brain regions functionally connected in the contrast motor > pseudo-verbs in red and mental > pseudo-verbs in green. Color bars show z scores. LH: left hemisphere; RH: right hemisphere.

Finally, psychophysiological interaction was calculated between categories of verbs. The motor > mental contrast did not show significant interaction, while the mental > motor contrast evidenced an interaction between the right cerebellum and medial occipital area (Fig 8, S8 Table). The occipital area corresponds to 8.2% of the medial occipital regions that showed an increase in BOLD signal in the mental > motor activation map resulting from the GLM analysis (Fig 5).

**Fig 8.**
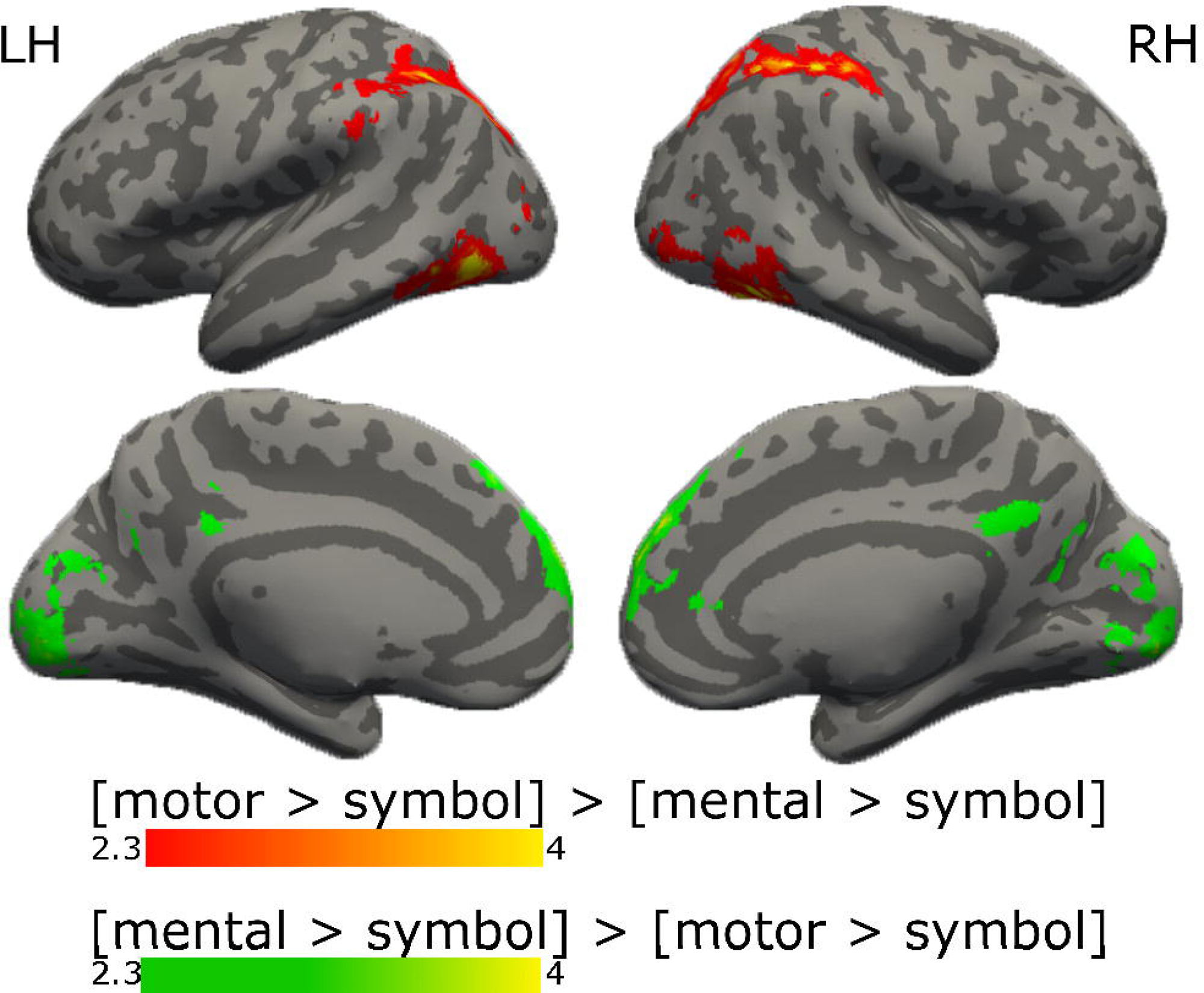
PPI results for the comparison between mental and motor verbs. Graphical representation of PPI results with the R-cerebellum seed, brain regions functionally connected in the contrast mental > motor green. Color bar shows z scores. Coordinate z slice in cerebellum = −29. LH: left hemisphere; RH: right hemisphere.

## Discussion

This study confirmed the differential recruitment and functional connectivity of neural regions according to the sub-lexical processing level for pseudo-verbs and lexical-semantic for verbs. Reading of verbs involved bilateral frontal, temporal, and parietal regions as described for the reading neural network (10). Pseudo-verbs, which represent the sub-lexical level of processing, recruited the left language network and the extrasylvian motor system, including the right cerebellum. Differences in neural activation between semantic categories showed that mental verbs, relative to motor verbs, were represented by greater activation of the left-language perisylvian network and extrasylvian regions, such as the cerebellum and occipital areas, which were functionally connected. According to our expectation, these results showed that visual modality-specific brain areas participate in processing mental verbs which convey abstract, non-motor actions, for example, “to understand”.

The results of the behavioral tests showed better linguistic performance for verbs and semantic fluency than phonological fluency. We take this to mean that the organization of the mental lexicon favors lexical-semantic representations over phonological characteristics. The verbal fluency of verbs was negatively correlated with the scores in the EHI, i.e., higher performance in the verb fluency task was related to less motor asymmetry. ROI approach, in the contrast mental versus motor verbs, detected that verbal fluency was positively correlated with the change in BOLD signal in the left lateral occipital cortex, which receives inputs from higher-order representations to distinguish and integrate meaning (45,46) for actions verbs and observed actions (8). Furthermore, lateralized motor behavior was associated with less signal change during reading mental verbs in semantic areas, i.e., left middle temporal gyrus, and motor areas, such as left SMA and right cerebellum, after eliminating the influence of verb fluency.

According to our expectation, compared to symbols and pseudo-verbs, verbs recruited an extended reading network on bilateral temporal regions that belong to the ventral stream, related to semantic processing (1,3) and medial regions related to attention (7,45). Verb reading did not recruit the dorsolateral prefrontal cortex since meaning selection is already established and stored in the mental lexicon. In contrast, pseudo-verbs reading required more support from the articulatory system, involving frontal regions of the left perisylvian circuit, which are part of the dorsal stream (3,10). In the present study, pseudo-verbs lack meaning but simulated verbs, i.e.,, since they were built with the expected ending of a verb in Spanish in the infinitive form (i.e.,, *ar, er, ir*). Relative to verbs, pseudo-verbs reading recruited the bilateral dorsolateral prefrontal cortex, involved in the search for meaning (12,45), the superior lateral occipital cortex, including the supramarginal gyrus, a semantic hub (20,46,47), and specific-modality systems such as the visual word form area (48), and motor regions (somatosensory cortex, SMA, and right cerebellum [crus I, crus II, lobules VI, and VIIIa]). However, compared to verbs, pseudo-verbs reading did not recruit semantic brain areas from the temporal lobe. These findings confirm a robust interhemispheric dependency for the semantic representation of verbs as part of the ventral stream; and a higher involvement of the left language network, dorsal stream (3), including sensory-motor areas for pseudo-verbs. Since motor areas have been associated with phonological and articulatory processing (10,17) and verbal working memory (49), future studies should untangle whether the recruitment of motor-specific regions can be attributed to semantic grounding on sensory-motor processes or to another type of processing during pseudo-verbs reading.

The comparison between semantic categories of verbs, i.e., motor to respect mental verbs, yielded a null result, which supports the notion that motor verbs, being concrete, require fewer neural mechanisms for semantic representation relative to abstract verbs (30). This finding was confirmed by the contrasts between each verb category and pseudo-verbs (data not shown). The only significant finding was found when the effect of visual processing from symbols was removed from the comparison ([mental>symbol] > [motor>symbol], supplementary material). In this contrast, the reading of motor verbs resulted in increased BOLD signal in bilateral posterior multimodal regions: superior lateral occipital cortex and ventral temporal regions involved in storing meanings and visual identification of objects (20,47).

On the other hand, mental verbs, compared to motor verbs, showed an increase in BOLD signal on the multimodal left language network–frontal and temporal regions–, on sensory-motor systems in visual medial occipital regions, and motor areas such as SMA and right cerebellum (crus I/II, and lobule VI). It is worth noting that the activation observed in the left language network and cerebellum did not survive when the effect of symbols was removed, and only the medial frontal and occipital regions maintained an increase in BOLD signal. When the effect of pseudo-verbs was removed, mental verbs recruited only visual systems from occipital regions. Our mental verbs were classified as abstract, less imaginable than motor verbs. Less imaginability could promote top-down modulation to increase visual perceptual processing for reading abstract verbs (50) and maybe use visual experiences to ground the semantic representation (8). The visual regions with increased BOLD signal also showed functional connectivity with the right cerebellum, suggesting that this may be a network that supports the semantic representation of abstract mental verbs. Thus, consistent with the grounded cognition framework, our results suggest that abstract concepts related to mental action are grounded in modality-specific brain systems typically engaged in visual perception (20) that interact with multimodal systems for semantic representation (17,51).

Left SMA and right cerebellum, in addition to the left-lateralized language network, showed an increased BOLD signal for both pseudo-verbs with respect to verbs, and mental respect to motor verbs, while the primary motor cortex was recruited only for pseudo-verbs. Since the SMA and cerebellum are not linked to motor representations like the primary motor cortex, they may participate more abstractly in the search for semantic representations. BOLD signal change while reading mental verbs in the SMA and cerebellum regions was related to lower motor asymmetry scores and functional connectivity with another cortical region. Thus, SMA and cerebellum may be grounding the abstract representations onto modality-specific sensorimotor mechanisms. This suggestion agrees with previous studies showing that the SMA, cerebellum, and medial posterior regions, including the precuneus, are nodes of the cognitive control network (52–54); the SMA has been associated with task switching (55), response inhibition (56), and action selection and planning based on internal goals (12,57,58). At the same time, the cerebellum is believed to create internal models of sensorimotor information for prediction and optimized execution (59–61).

As already mentioned, performance in verbs fluency and lateralized motor activities (i.e., EHI scores) were related to the change in BOLD signal associated with the semantic contrast (mental vs. motor verbs) on four regions of interest determined with an automated meta-analysis for ‘verbs.’ Two of those regions are semantic regions, i.e., the left superior lateral occipital cortex and the left middle temporal gyrus. The others are motor regions, i.e., left SMA and right cerebellum. Firstly, functional connectivity for each semantic category of verbs and pseudo-verbs with respect to symbols was obtained. Left superior lateral occipital cortex, as seed, showed functional connectivity with bilateral frontal, parietal, and temporal regions for both categories of verbs, while for pseudo-verbs, it was coupled only with the left ventral frontal region, including primary sensorimotor regions. It has been suggested that the left superior lateral occipital cortex functions as a semantic convergence zone (45,62) that could manage the meaning of stimuli. For verbs, the left superior lateral occipital cortex was coupled with those regions that store conceptual knowledge, which are quite widely distributed. For pseudo-verbs, this region and left temporal middle gyrus showed functional connectivity with frontal regions involved in searching for meaning and learning (12,63), somatosensory representations from pre and postcentral gyri (63), articulatory and motor processing (10).

Interestingly, the motor seeds (left SMA, and right cerebellum) showed functional connectivity only for mental verbs compared to symbols. Left SMA was coupled with anterior regions, while right cerebellum with bilateral posterior regions such as temporal, medial occipital, sensorimotor regions, and polar frontal. These findings strengthen the idea that SMA and cerebellum collaborate to ground the abstract to modality-specific representations and support previous studies which have reported that conceptual meaning at the abstract level is embodied by the interaction of sensory and motor mechanisms with multimodal processing (51). This interaction collects a variety of sparse bodily experiences (31,33) or strategies (8,10) for semantic representation.

Secondly, functional connectivity for each category of verbs compared to pseudo-verbs was calculated for four seeds. Again, the superior lateral occipital cortex was functionally connected with an extended bilateral temporal and parietal network. Furthermore, the left SMA showed functional connectivity, for both motor and mental verbs, with bilateral superior temporal regions, posterior cingulate cortex, and frontal pole. Similarly, the right cerebellum showed functional connectivity with other regions only for mental verbs, including bilateral temporal regions, post and precentral gyri, frontal areas, and precuneus. Finally, the connectivity analysis demonstrated that a portion (8.2%) of medial visual areas recruited by reading mental verbs compared to motor verbs were functionally coupled with the right cerebellum in the same contrast, i.e., mental verbs versus motor verbs.

Our seed-ROI in the right cerebellum was in crus I and lobule VI. The involvement of the cerebellum in language functions is strongly lateralized to the right hemisphere (14–16), specifically lobules VI, VII, and crus I/II have shown functional connectivity with the cortical language network, and support for semantic prediction in speech production and comprehension (64,65) beyond articulation as a motor component of language (54,60). It has been proposed that the cerebellum creates internal models of sensory-motor information and updates them based on the comparison with the actual outcome (59–61). Thus, it is possible that during reading pseudo-verbs, and mental compared to motor verbs, the cerebellum provided the internal models of sensory-motor information to ground the representations or mapped the semantic representations when the referent was lacking (21). Likewise, the right cerebellum was coupled with the visual cortex when reading mental verbs, supporting the idea that the cerebellum creates sensory-motor internal models connected with the visual sensory system to optimize language comprehension (59–61,66). Recently, a study has shown that crus II and lobule VI in cerebellum are spontaneously and functionally coupled to the primary visual cortex (67). Previous reports have involved both cerebellar areas with predictive coding on visual areas (50,64,65). Our results strengthen the idea that the cerebellum is responsive to semantic information, and that it is part of the language network that extends beyond the left language network. More studies are needed to reveal what modulation is exerted from the cerebellum to cortical areas during language processing. Knowing how the cerebellum modulates the flow of information between specific-modality systems in cortical areas will help explain its more significant involvement in abstract versus concrete concepts.

The current study’s limitations are that other lexical levels, such as nouns or adjectives, were not included. Pseudo-verbs were built following the phonological rules of Spanish. However, we did not test their concreteness-abstraction, imageability, or arousal, which could be very useful in comparing them to semantic categories of verbs. Since it has been reported that different types of abstract concepts are associated with representational differences (17,18), adding different categories of abstract verbs could have been useful.

In conclusion, this study confirmed differential activation and functional connectivity for reading verbs and pseudo-verbs. According to the dual stream model, the left dorsal stream that supports the sub-lexical route, and the extrasylvian motor system, including the right cerebellum, were involved in reading pseudo-verbs; while the ventral stream that maps words onto lexical conceptual representations was involved in reading verbs. Our findings support the embodied or grounded cognition model in that modality-specific brain regions contribute to the semantic representation of abstract verbs and the well-established and multimodal left perisylvian networks. Furthermore, a preferential modality-specific system was detected: visual systems were recruited by abstract verbs and showed functional connectivity with the right cerebellum forming a network that supports the semantic representation of abstract concepts. These results confirm the dissociation between sub-lexical and lexical-semantic processing and provide evidence for the neurobiological basis of semantic representations grounded in modality-specific systems for abstract concepts.

## Supporting information

https://docs.google.com/document/d/1kpWejNf3ZWQxAFLUfwteyJeWaaeKGbBB/edit?usp=sharing&ouid=114306660564224461935&rtpof=true&sd=true

## Notes

### Competing Interest Statement

The authors have declared no competing interest.

