## Supplementary material for "Contribution and functional connectivity between cerebrum and cerebellum on sub-lexical and lexical-semantic processing of verbs": https://docs.google.com/document/d/1kpWejNf3ZWQxAFLUfwteyJeWaaeKGbBB/edit?usp=sharing&ouid=114306660564224461935&rtpof=true&sd=true

**RESULTS**

fMRI results

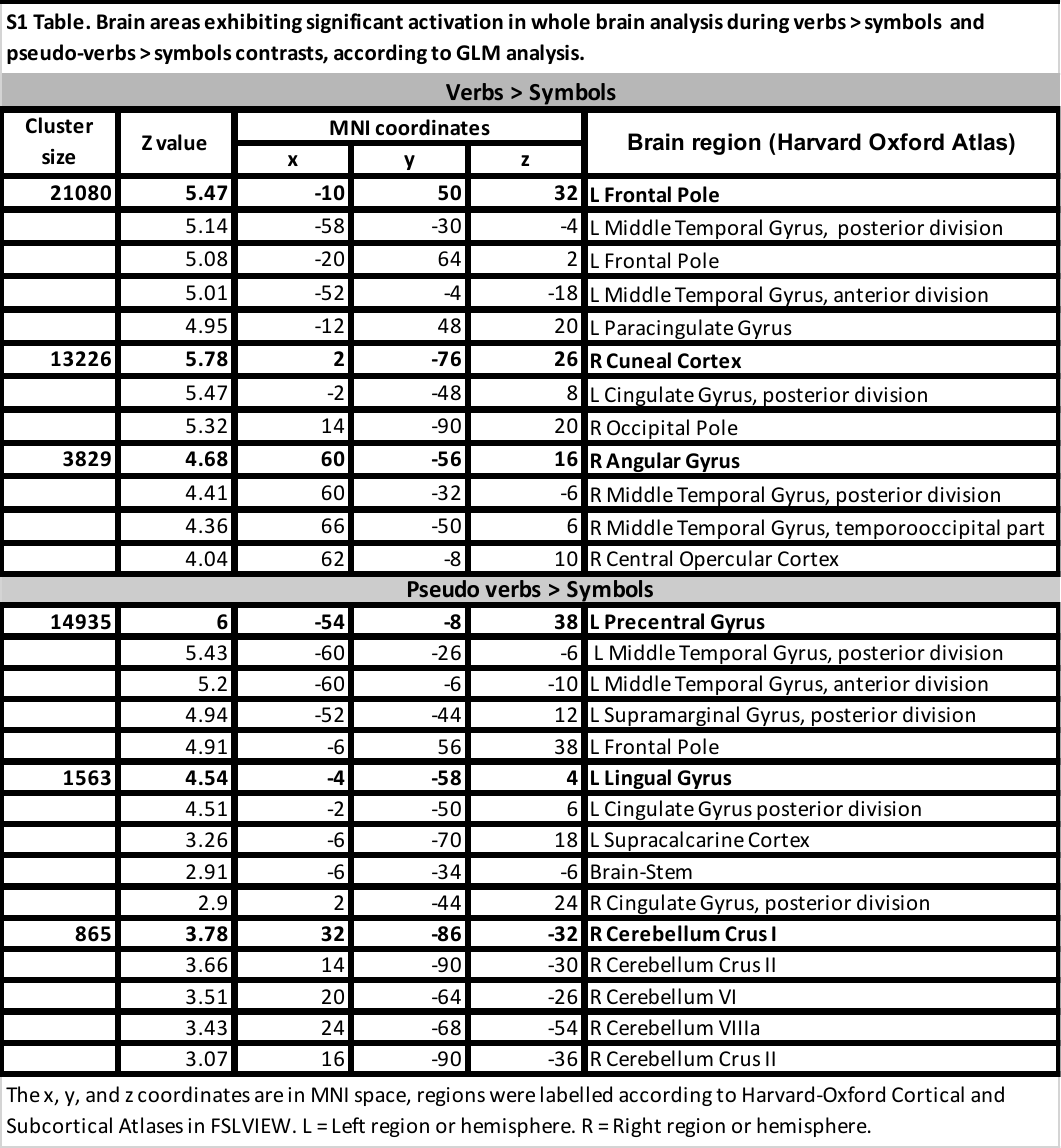

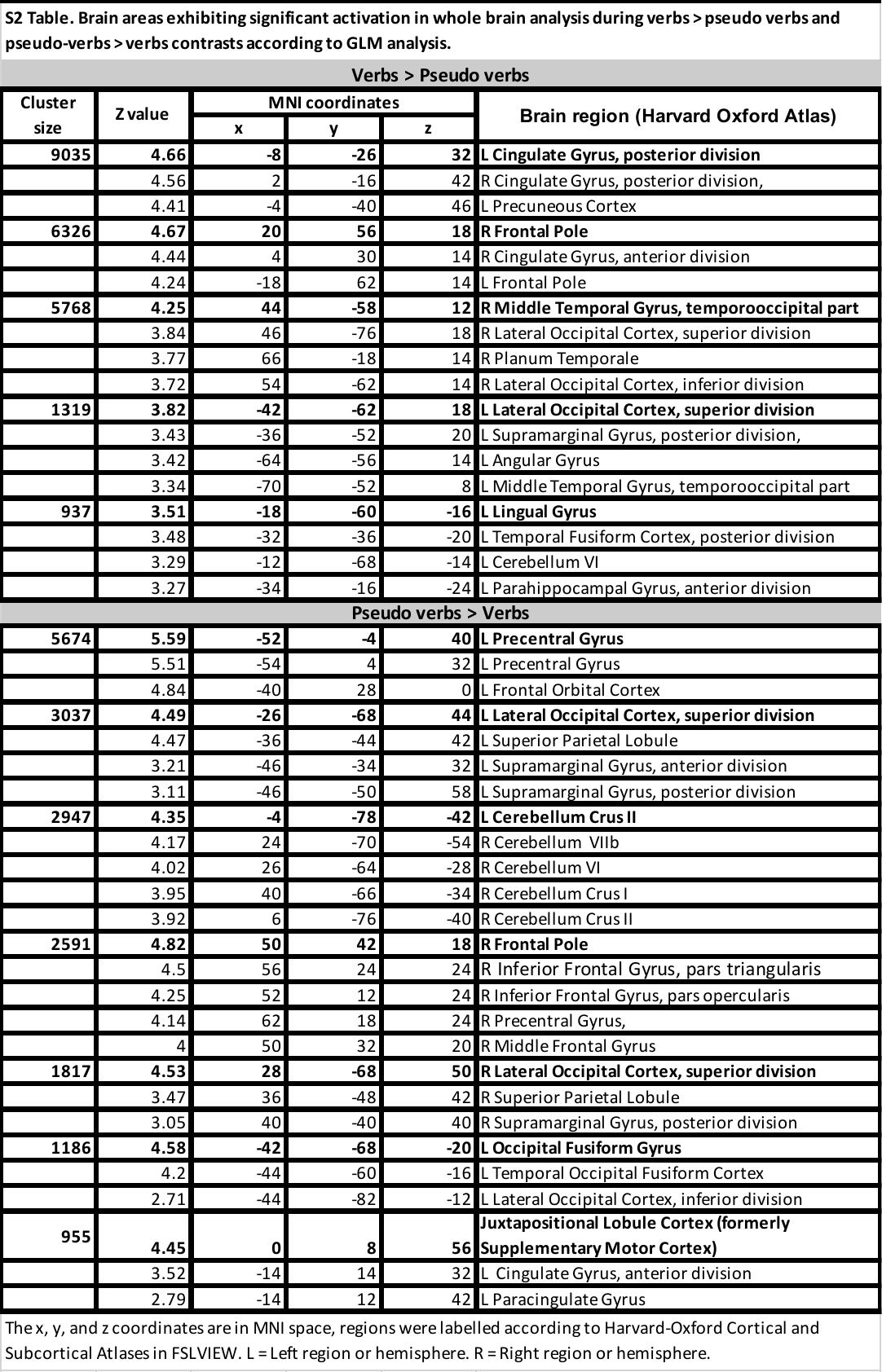

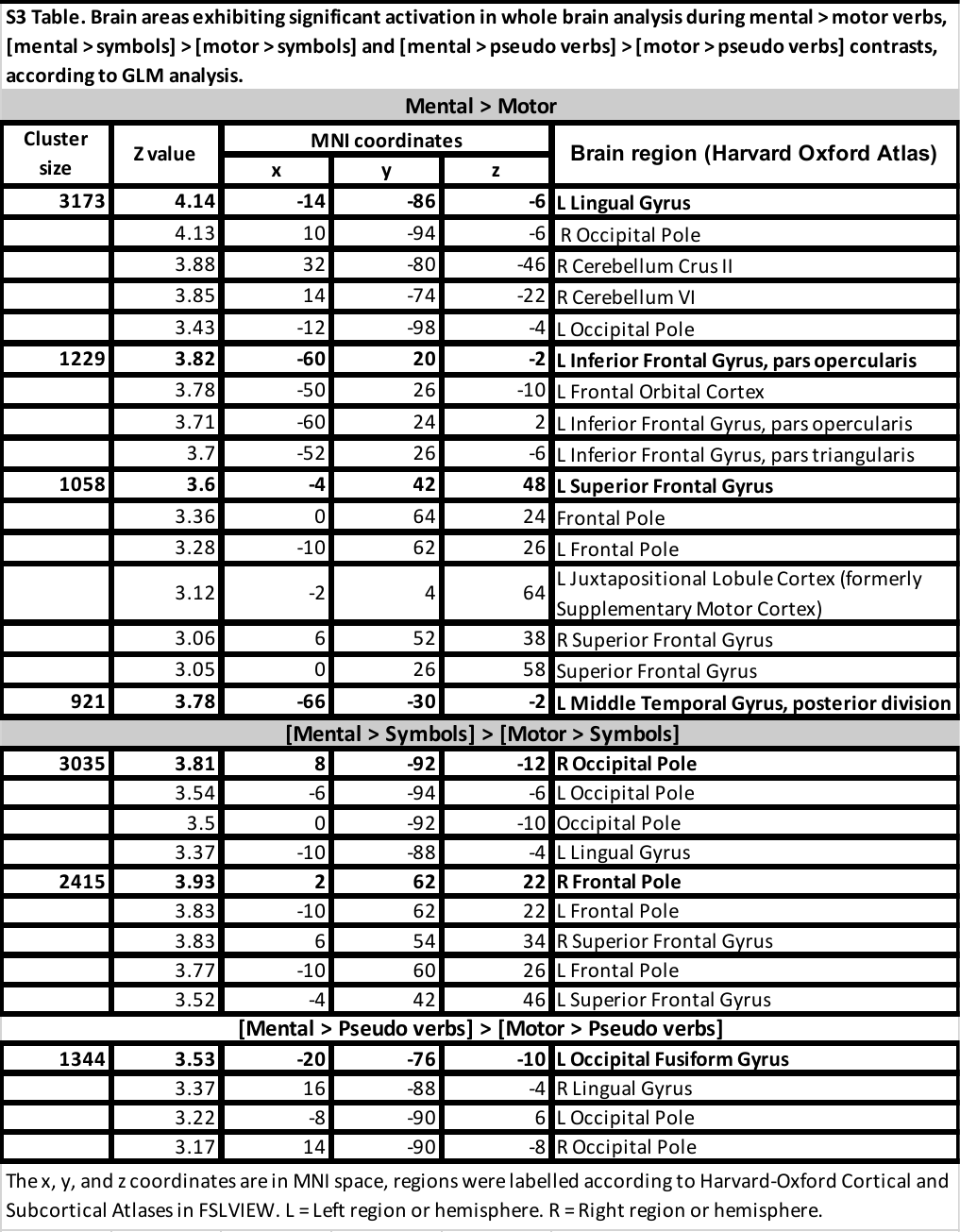

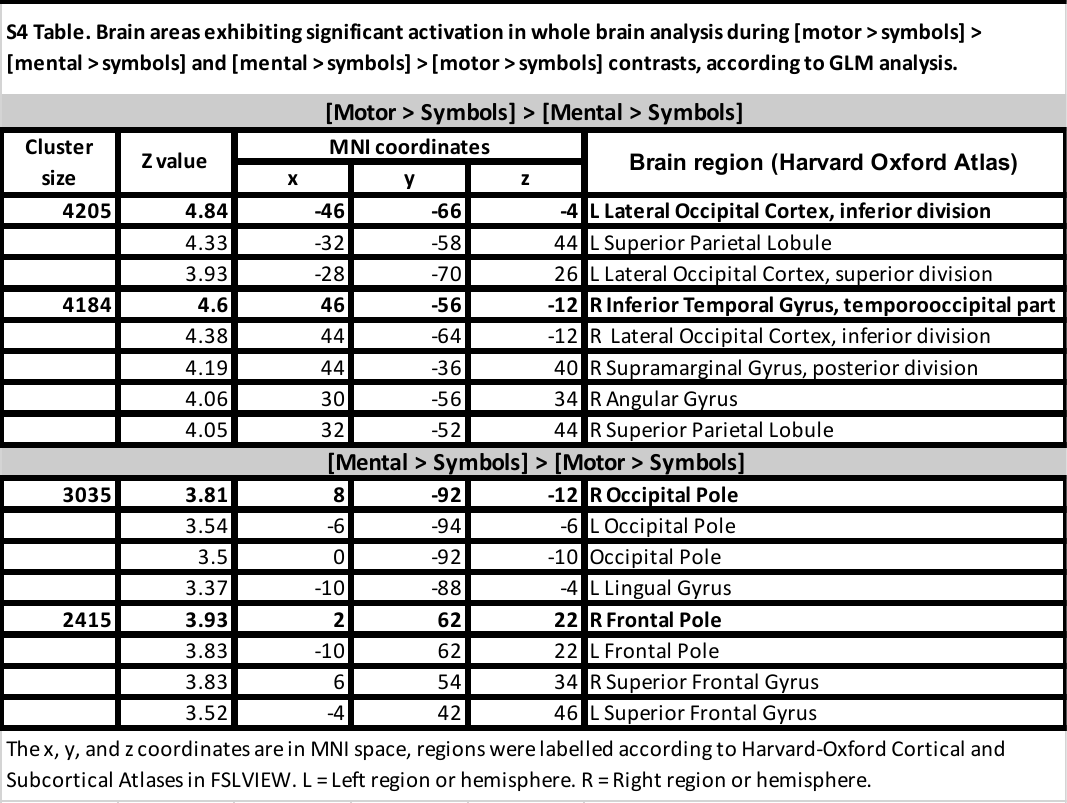

**S1 Fig**. **Activation maps for the** **comparison between motor and mental verbs when removing the effect of visual processing (i.e., symbols)**. Graphical representation of GLM's results, brain regions activated in the contrast [motor > symbol] > [mental > symbol] in red, and [mental > symbol] > [motor>symbol] in green. No other contrast between motor compared to mental verbs showed significant differences. Color bars show z scores. LH: left hemisphere; RH: right hemisphere.

Region of interest analysis

We then asked how this BOLD signal change during verbs categories was related with specific brain areas related to language processing. For this, ten ROIs were defined from meta-analysis for 'verbs' (see methods).

In addition to the correlations reported in the main text, we detected the following FWE corrected correlations. The BOLD signal change in L-SMG ROI for verbs > symbol contrast was related to phonological fluency (r = -0.56, p < 0.01). While, for verbs > pseudo-verbs contrast, R-STG ROI was related to semantic fluency (r = -0.59, p < 0.01), and verbs fluency was related to L-LOC (r = -0.53, p < 0.01) and L-SMG ROIs (r = -0.59, p < 0.01) when we calculated a partial correlation with phonological fluency and EHI, respectively. All these negative correlations were corrected to p < 0.01.

**S5 Table. Correlation between signal BOLD and behavioral performance.**

| **Contrast** | **Brain región** | **Behavioral task** | **Partial correlation** | **r** | **p <**  ***corrected** |
| --- | --- | --- | --- | --- | --- |
| verbs > symbol | L-SMG | phonological fluency | NA | -0.56 | 0.01* |
| verbs > pseudoverbs | R-STG | semantic fluency | NA | -0.59 | 0.01* |
|  | L-LOC | verbs fluency | phonological fluency | -0.53 | 0.01* |
|  | L-SMG | verbs fluency | EHI | -0.59 | 0.01* |
| mental > motor verbs | L-MTG | EHI | NA | -0.54 | 0.01* |
|  | L-LOC | verbs fluency | NA | 0.53 | 0.01* |
|  | L-SMA | EHI | verbs fluency | -0.45 | 0.05 |
|  | R-Cerebellum | EHI | verbs fluency | -0.42 | 0.05 |

L-SMG = left supramarginal gyrus (-60, -26, 28), R-STG = right superior temporal gyrus (56, -32, 2), L-LOC = left superior lateral occipital cortex (-26, -60, 48), L-MTG = left posterior middle temporal gyrus (-58, -10, -14), L-SMA = left supplementary motor area (-2, 8, 60), R-Cerebellum = right cerebellum (34, -64. -29), EHI = Edinburgh Handedness Inventory, NA = Not Applicable, * = FWE corrected.

PPI results

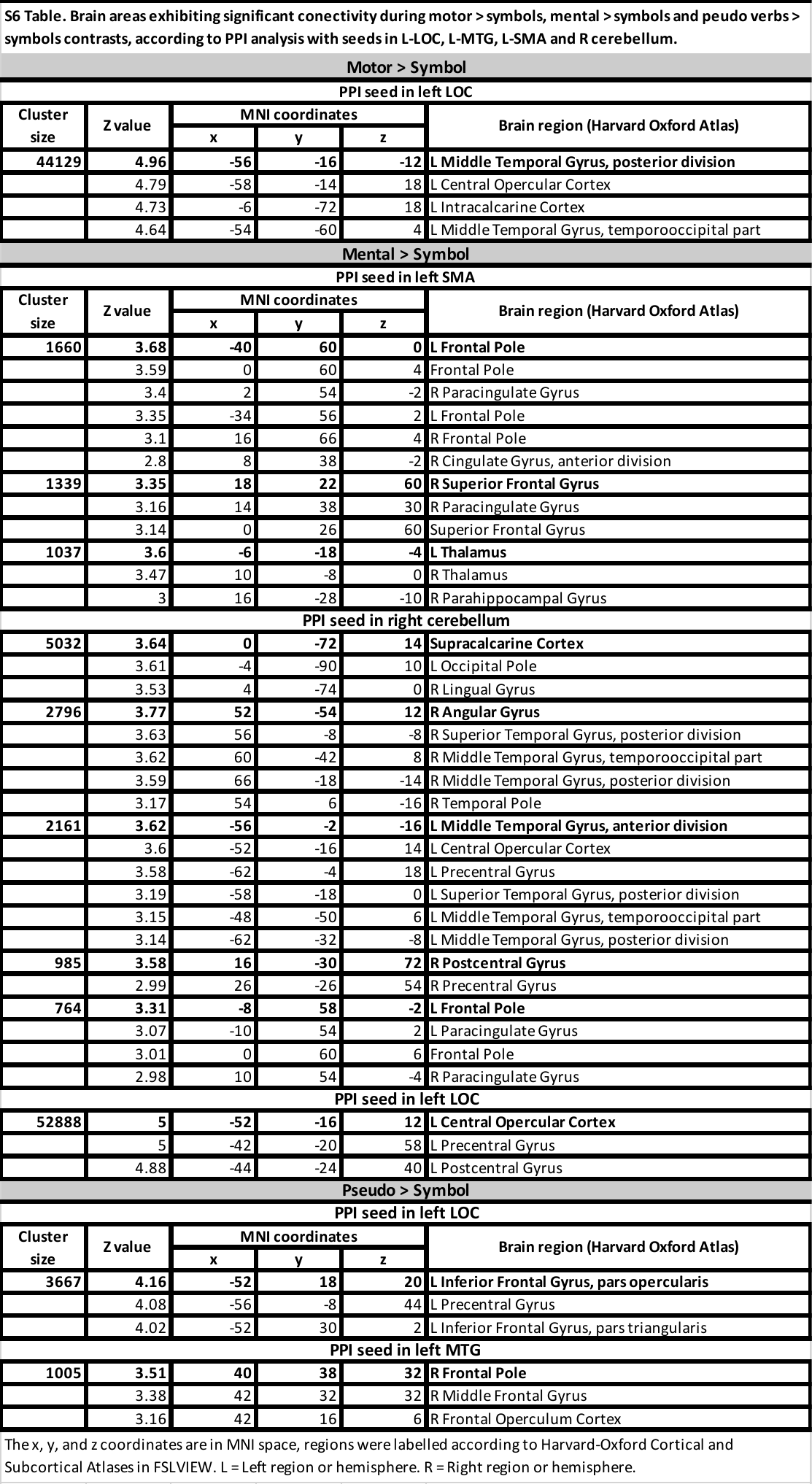

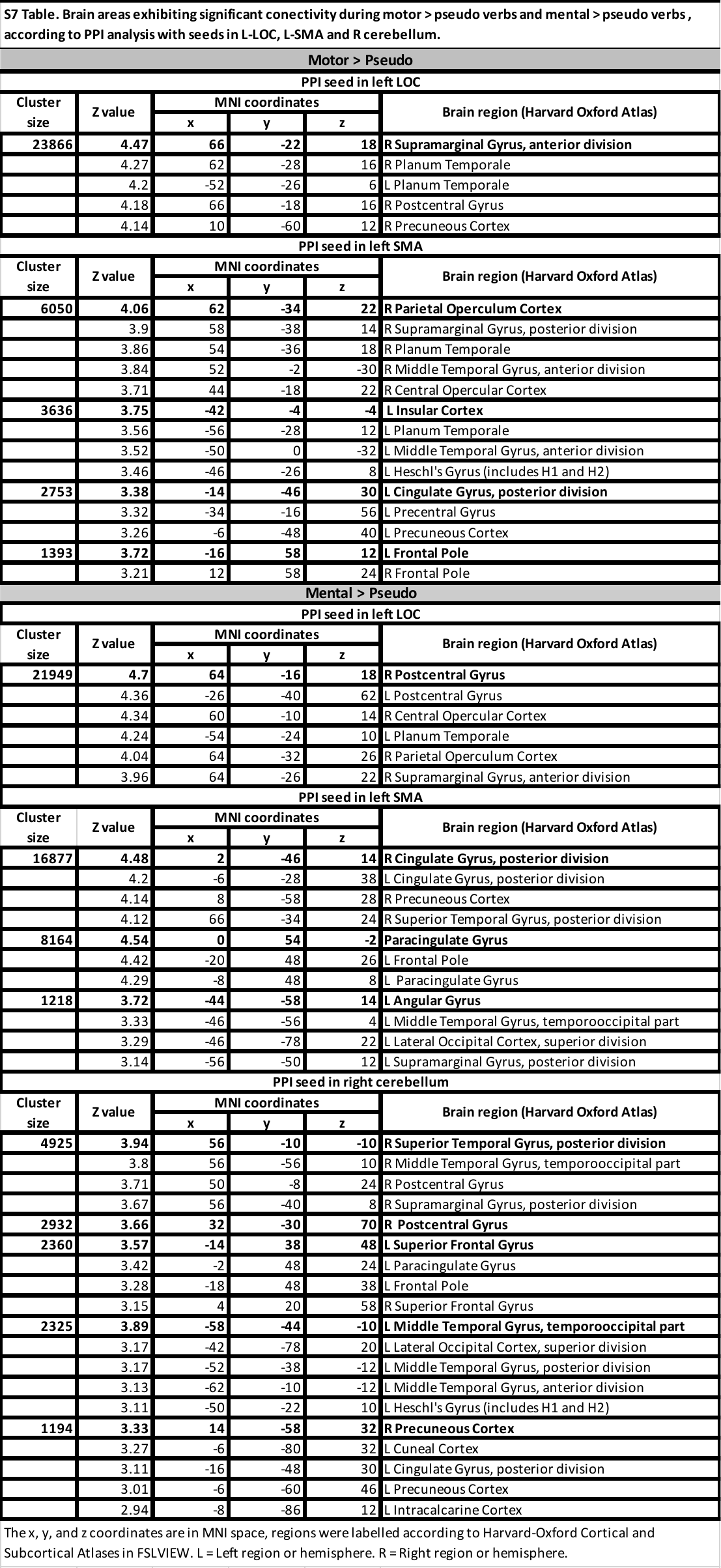

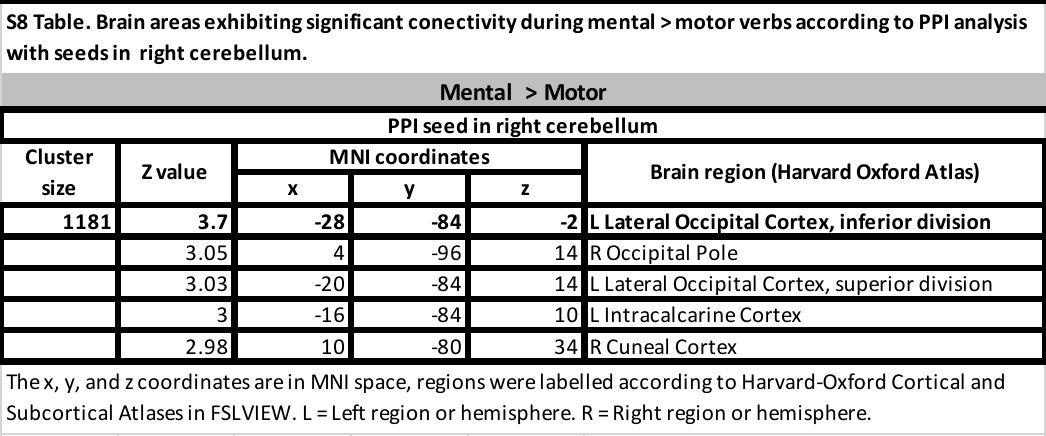
